# Disentangling the contributions of stress fibres and the unbundled actin meshwork to the anisotropy of cortical tension in response to cell shape

**DOI:** 10.1101/2024.10.25.620349

**Authors:** F. Fage, S. Asnacios, A. Pluta, A. Richert, C. Vias, J. Fouchard, H. Enslen, J. Etienne, A. Callan-Jones, M. Thery, D. Pereira, A. Asnacios

## Abstract

Many fundamental biological processes, in particular development and morphogenetic movements, involve tissue and cell deformation, as well as the generation of anisotropic mechanical stresses. They are often accompanied by the appearance of oriented contractile actomyosin structures resembling the stress fibres (SF) observed *in vitro*. Here, we investigate, at the single cell level, how cell shape — by itself — could control the structure and tension of the actomyosin cortex. Using a unique combination of 3D micropatterning, single peripheral SF (PSF) tension measurement, laser ablation and image analysis, we show that cell shape anisotropy, e.g. its 2D aspect ratio, is indeed sufficient to induce anisotropy of the cortical structure and tension. In particular, taking into account the experimentally measured anisotropy of the cortical meshwork, we could quantify cortical tension and decouple the contribution originating from bundled actin (oriented cortical stress fibres, CSF) and the contribution of the unbundled actin meshwork (UAM). We show that the increase of cortical tension anisotropy with the cell’s aspect ratio depends on the CSF alignment and orientation, the contribution of the isotropic mesh being independent of cell shape. Remarkably, while experimental data from single stress fibre measurements and laser ablation were analysed through different theoretical frameworks, namely that of negative pressure in nematics and hole drilling in prestressed materials, we found quantitatively the same composite material behaviour. In sum, we decipher here the very material properties of the actomyosin cortex, and its sensitivity to cell shape which is at the root of many mechanobiological processes, in particular morphogenesis.

## I. INTRODUCTION

Contractile actomyosin fibres are often observed in living organisms, in particular during development, where they seem for instance to delimit different segments of the body and to follow specific symmetries [1]. The question then arises whether these fibres lead to the observed shapes and symmetries, or conversely whether the developing shapes, by themselves, induce specific spatial distributions and orientations of contractile fibres that could then correctly structure the developing body. Answering this fundamental question in developing organisms is quite challenging, in particular due to the interplay between the genetic programs involved in shape establishment [2], and the feedback of the physical and mechanical cues arising from the tissue shape itself [3]. Indeed, recent observations suggest that actin fibres, their orientation and even topological defects in their nematic order could control the organisation of cell assemblies [4] and the morphogenesis of hydra [5].

One way to improve our understanding of such complex processes is to consider single cell systems where one can investigate the effect of shape on the distribution and orientation of contractile fibres without the interference of developmental genetic programs. Thus, we decided to study, in vitro, whether imposing cell shape, and more precisely its aspect ratio in 2D, could induce specific ordering of the actomyosin cortex with well-defined spatial distributions of stress fibres (SF), and whether this could affect its surface tension. In other words, we asked the simple question: does cell shape, by itself, impact the structure and material properties of the cell cortex?

The actin cortex is a thin 2D crosslinked biopolymer network situated beneath the plasma membrane of animal cells. It is recognized to control cell shape through non-muscle myosin II powered contractility [6]. Actomyosin contraction generates cortical tension, a cell surface tension, *i*.*e*. a force per unit length, which compensates for pressure difference across the plasma membrane due to osmotic pressure [7]. Cortical tension can either be isotropic and uniform throughout the cortex for instance when cells round up during mitosis [8], or display anisotropy and/or gradients with more contractile regions leading to shape changes, morphogenesis [2], like in cytokinesis [9], or more generally in development [10]. Thus, understanding the relationship between the structure of the cortex and the spatial regulation of cortical tension is fundamental. Cortical stress fibres (CSF), which are bundled contractile actomyosin fibres embedded in the cell cortex, act as force dipoles generating oriented contraction [11,12] at the cell scale [13]. Thus, the orientation of CSF could potentially control the anisotropy of cortical tension, in particular in response to cell shape [14]. However, while the cortex architecture is demonstrated to regulate cortical tension [15], measuring tension and the specific contribution of CSF is challenging, in particular because the unbundled cell cortex and the embedded CSF constitute a mechanical continuum [16].

Fortunately, with the emergence of micro-patterning, it has been shown that non adhesive regions at the cell periphery always produce peripheral stress fibres (PSF) [17], the mechanics and geometry of which have turned out to be valuable reporters of cortical homeostasis [16,18–20]. Indeed, since PSF are mechanically linked to the cortex, their radius of curvature is set by a balance between PSF internal tension (line tension acting along the PSF, pulling them straight) and the cortical tension acting laterally on the PSF (surface tension that tends to curve the PSF). However, beyond cortical tension, many other parameters, such as the cell spread area [21,22], the cell’s total free (non-adherent) perimeter [17], the free non-adherent length [18], the structural anisotropy [20,23], the rigidity of the substrate [24] or the detailed local angular distribution and connections of CSF [19], have all been shown to influence peripheral SF features, leading to different models and physical representations of the cell cortex [14,24,25]. In order to determine the specific effect of cell shape anisotropy on cortical structure and tension, we first designed original micropatterns allowing us to distinguish the effect of the free length of PSF from that of the overall cell shape, while keeping constant the cell area and the total free perimeter. Then, we designed specific cantilevers shaped and calibrated to allow us to measure tension along each side of the cell. Thus, we were able to quantify, in each individual cell, the anisotropy of cortical tension and its evolution with cell aspect ratio. This cell shape-induced anisotropy of cortical tension was further confirmed by laser ablation. Ultimately, combining laser ablation with analysis of SF orientations and modelling the cell cortex as a material composed of oriented force dipoles embedded in an isotropic contractile meshwork, we could reveal the specific contribution of SF to the anisotropy of cortical tension, and its dependence on cell shape.

## II. MATERIAL AND METHODS

### A. Cell culture

LifeAct Fared fluorescent 3t3 cell lines were obtained from stable piggyback transfection and single colony selection. Special care has been taken concerning the proliferation, spreading and fluorescent signal within the selected colony. They were grown in DMEM supplemented with 10% FBS (Eurobio) and 1% Penicillin-Streptomycin. They were cultured at 37°C with 5% CO2.

### B. Plasmid construction

The Blasticidine-CAG-LifeAct-FarRed plasmid was constructed using a piggybac plasmid: Bsd-CAG-MCS as a backbone. This backbone has been digested by the BamHI restriction site following a standard digestion protocol. The enzyme used was purchased from New England Biolabs (NEB, Beverly, MA). The gene FarRed is PCR amplified from pmiRFP670-N1 plasmid (Addgene plasmid 79987) and the peptide LifeAct is PCR amplified from an internal plasmid. Each amplicon makes respectively 963 bp and 117 bp. The PCR cycling parameters were: 98°C for 3 min, 5 cycles of 98°C for 20 s, 65°C for 20 s, and 72°C for 30s, 20 cycles of 98°C for 20 s and 65°C for 30 s, 98°C for 20 s, 83°C for 20 s and 72°C for 30s, and 1 cycle of 72°C for 5 min. The purified vector and inserts mixture were blended with 15 μL Gibson assembly homemade mix and incubated at 50°C for 20 min.

Then, 50 ng of Gibson reaction was used to transform 200 μL of home-made Stbl3 competent cells following a standard transformation protocol. The mini preparations of plasmid DNAs were carried out using NucleoSpin Plasmid DNA purification kit (from Macherey-Nagel) and positives clones were sequenced.

### C. Fluorescent staining of fibronectin and cell nuclei

Nuclei were tagged in DAPI by incubation with 0.5 mg/ml Hoechst 33342 (Thermofischer) for 30 minutes followed by rinsing with DPBS and adding fresh medium. Fibronectin was tagged in RFP with the fluorescent labelling kit Ab Cy2 Labelling kit (GE Healthcare).

### D. 2D and 3D substrates preparation

The substrates were made using soflithography techniques. A silicon wafer was spin-coated with negative photoresist (SU-8 2005 Microchem Corp.) to obtain a thickness of 5μm. During UV insolation step, a chromium mask with the desired shape was used. For 3-D substrates, poly(dimethylsiloxane) (PDMS) (Sylgard 184; Dow Corning) was spin-coated on the SU8 mold to obtain a thickness of 100 μm, then cured at 65°C overnight. The thin layer was peeled off and placed gently in the bottom of a glass petri dish, with the 3-D microstructures facing-up. For 2-D substrates, PDMS was poured on the SU8 mold to get a layer of 1 cm and then cured at 65°C overnight. This thick PDMS layer was used as a stamp for the 2-D substrates. In addition, a glass coverslip (22x22mm) was covered with a thin layer of PDMS using a spincoater in a two-step process (first step at 2000 rpm for 30s, second step at 6000 rpm for 1min30s) to get a 10μm layer. This coverslip covered with PDMS will be use as substrate for the 2-D micropatterns during the stamping step. 2-D and 3-D substrates were placed in a UV-ozone cleaner (UVO-Cleaner®) during 15 min in order to clean and prepare the surfaces for stamping. At the same, the thick PDMS layer and a flat PDMS stamp were covered with fibronectin at 50 μg/ml for 45 min. Right after the cleaning step, 2-D (coverslip with PDMS) and 3-D (microstructures) substrates were stamped using the thick PDMS layer and the flat PDMS stamp, respectively (for details see [26]).. Finally, the 2-D and 3-D substrates were passivated with 1% Pluronic® F-127 (Sigma) for 45 min and rinse with PBS (1X). Substrates were stored for up to one week in PBS at 4°c.

### E. Line tension measurements of peripheral stress fibres

Line tension measurements were performed using a subcellular microrod probe based on the experimental setup previously described [27]. Cells were seeded, for 6 to 8 hours, on the 3-D fibronectin-coated-microstrutures and placed under the microscope (for details see section “Live imaging”). Measurements were performed on 3-D microstructures displaying an unique and fully spread cell (i.e. unique nucleus, based on nucleus labelling (DAPI)). The 2 μm diameter glass-microrod probe was put in contact with the peripheral stress fibre (PSF), and then pushed again the PSF at 1μm/s for 5s. The position of the tip was imaged at 10 fps and extracted using imageJ plugin Trackmate. The line tension is retrieved from the position of the tip, the displacement of the microrod base, the stiffness of the microrod, the curvature and the length of the peripheral arc andthe projection of the force applied by the microrod to the stress fiber.

### F. Live imaging for line tension measurements

Fluorescent live-cell images were obtained using an Olympus IX71 inverted epifluorescence microscope equipped with an Olympus UPLSAPO 60x W/1.2 NA objective, an orca flash 4.0 CCD camera (Hamamatsu), an X-Cite ® exacte fluorescence lamp (Excelitas Technologies), and a temperature controller (Air-Therm, World Precision Instruments). Vibration isolation was achieved by a TS-150 active antivibration table (HWL Scientific Instruments Gmbh). (Life Imaging Sciences). FluoroBrite DMEM Media cell culture media (ThermoFischer Scientific) supplemented with 10% fetal bovine serum, 1% penicilin/streptomycin and 25mM Temperature inside the chamber was kept at 37°c.

### G. Microrod manufacture, calibration, cleaning and coating

The microrod was obtained by stretching a 1 mm diameter glass rod with a micropipette puller (P1000, Sutter Instrument). A microplate with a known stiffness was used to calibrate the microrod along the two orthogonal axis used for line tension measurement (labelled x and y in figure 2). We measured a stiffness of 10.6 nN/μm and 7.7 nN/μm for the x and y axess, respectively. Prior to each experiment, this microrod was cleaned 30 min in a “piranha” mixture (67% sulfuric acid + 33% hydrogenperoxyde) and leaved in water overnight and then coated, for 45 min, with 1% Pluronic® F-1275 (Sigma) to prevent microrode-cell adhesion.

### H. Laser ablation

Photoablation was performed on an Eclipse Ti-E Fully-Integrated, Motorized Inverted Microscope system (Nikon) and using the iLas3 device (Gataca Systems) equipped with a passively Q-switched laser (STV-E, ReamPhotonics, France) at 355 nm producing 500 picoseconds pulses. Laser displacement, exposure time and repetition rate were controlled via ILas software interfaced with MetaMorph (MetaImaging). Laser photoablation and subsequent imaging were performed with 100x NA 1.49 oil objective azimuthal TIRF (Olympus). Temperature inside the chamber was kept at 37°c and CO2 at 5 %. Prior to laser ablation, cells were grown for 6 hours on 2-D micropatterns coated with fibronectin. On fully spread cells, a 5 μm diameter disc of cortical actin ablation was performed during live acquisition. For all photoablations, 25 repetitions of 25 ms pulses were used with 60 % of the 355nm laser power, corresponding to a pulse of approximately 1s.

### I. Analysis of the stress fibres orientations

Stress fibres orientations were measured only over the cortical stress fibres, the peripheral stress fibres (i.e. the peripheral arcs) being excluded from the measurement. The orientations of the cortical stress fibres were determined using OrientationJ Distribution plugin (Rezakhaniha et al., 2012) in ImageJ (NIH). The order parameter of the actin cytoskeleton, for each cell, was calculated as ⟨*S*⟩ = ⟨cos 2*α*⟩, where *α* is the angle with the mean orientation (i.e. the nematic director *n*) (Appendix C).

The structure tensor for each pixel was calculated using a Gaussian analysis window of size 10 pixels and a cubic spline gradient. From the component Aij of the structure tensor and its eigenvalues and, the directional orientation 1/2arctan(2(Axy/Ayy - Axx)), the energy Axx+Ayy and the coherency (λmax λmin)(λmax+λmin) were calculated for each and a histogram over all pixels was calculated. Only pixels with energy larger than 10% of the maximum energy were considered in the histogram to reduce fluorescence noise. In addition, the histogram was weighted by the coherency value to fit at best anisotropic structures such as SFs. Only pixels with a coherency larger than 10 % were considered.

## III. RESULTS

### A. Global cell shape controls the radius of curvature of peripheral stress fibres

When cells are spread on adhesive micropatterns, PSF form on cell edges, over non-adhesive regions. Since the actin cortex is mechanically connected to PSF [14,18,19], the morphology of these fibres is sensitive to the traction exerted by the cell cortex. Thus, we first measured the radii of PSF in micro-patterns of different aspect ratios through image analysis. However, since the radius of PSF is known to depend on the free length between adhesive regions [18], we designed original 3D micropatterns allowing us to distinguish the effect of the free length of PSF from that of the overall cell shape, while keeping constant the cell area and the total free perimeter (Fig. 1, and SM section 1). We made rectangular micropatterns with aspect ratios varying from 1 to 3 (defined as the ratios between the lengths of the long and short sides of the micropatterns). Independently from their aspect ratio, all micropatterns displayed the same free length along the short and long axes of the cells: d_S_= d_L_= d_1_=15 μm (Fig. 1a and 1b). In sum, thanks to the original design of our micropatterns, we could vary the cell aspect ratio without varying the free length of the PSF, allowing us to determine the specific effect of cell shape anisotropy on cortical tension and structure.

**FIG 1.**
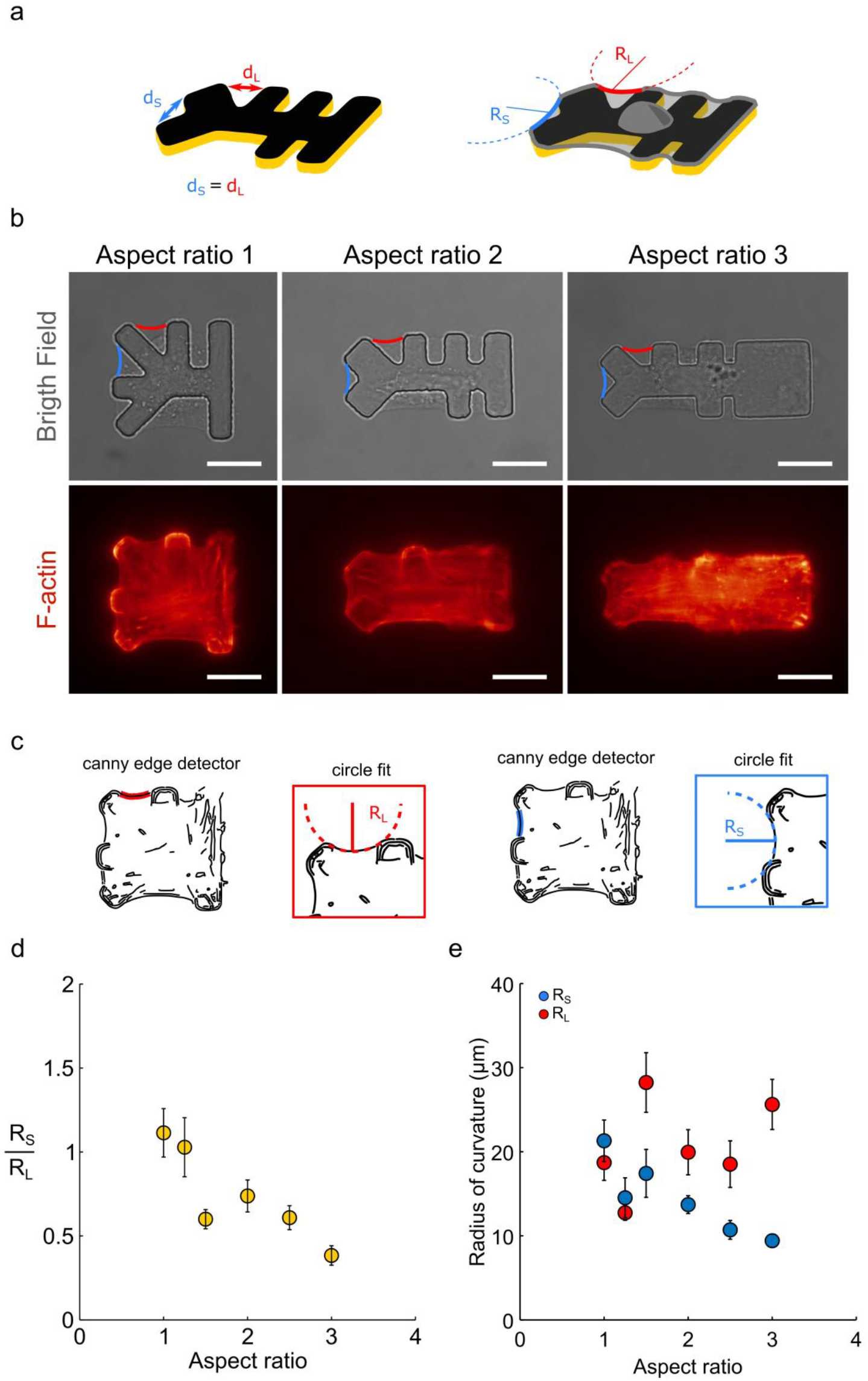
(a) Schematic representation of a 3D micropattern with an aspect ratio of 2 and two free lengths *d*_*S*_ (blue) and *d*_*L*_ (red) both measuring 15 μm, but situated along orthogonal sides of the micropattern (left). Cells spread on this kind of micropatterns display two peripheral stress fibres overs these free lengths, with radii of curvature *R*_*S*_ (blue) and *R*_*L*_(red) respectively (reight). (b) Bright field and fluorescence (F-actin, red hot) images of cells spread on micropatterns of 2000 μm^2^ with an aspect ratio ranging from 1 to 3. (c) Procedure of arc radius fitting. First, the cell contours are segmented with a canny edge detector filter. Then, both ends of each stress fibre are manually determined and a circle is fitted to the arc fibre using Pratt’s Method. (d) Radii of curvature *R*_*s*_ (blue) and *R*_*L*_ (red) as function of cell aspect ratio. (e) Cell-by-cell ratio 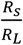 of radii of curvature as a function of cell anisotropy. (number n of cells n_AR1_=21, n_AR1.25_=17, n_AR1. 5_=15, n_AR2_=16, n_AR2.5_=13, n_AR3_=15, number N of independent experiments N_AR1_=2, N_AR1.25_=1, N_AR1. 5_=3, N_AR2_=1, N_AR2.5_=2, N_AR3_=2).

Measuring the curvature radii through image analysis and circle fits (Fig. 1c and), we found that increasing the cell aspect ratio (AR) has no significant effect on the curvature radius R_L_ of the fibres generated along the long axis of the cells, with R_L_ ∼ 20 μm regardless of the AR, whereas the radius R_s_ along the short axis decreases from ∼20 to ∼10 μm for increasing AR (Fig. 1d). This cell-axis dependent behaviour is even more striking when one considers the cell-by-cell ratio R_S_/R_L_ of the PSF radii, its mean value being a clearly decreasing function of the AR (Fig. 1e). This demonstrates that a global parameter (cell shape anisotropy) can control a local feature of PSF (their radius of curvature), and led us to hypothesize that cell shape anisotropy induces an anisotropy of the cortical (surface) tension pulling perpendicularly on PSF. Along this line, we observed that PSF situated on opposite sides of the cells displayed similar fluctuations of their curvature suggesting a correlation at distance through cortical tension (SM section 1). Thus, we decided to determine the dependence of cortical tension on cell shape anisotropy.

### B. Line tension of PSF is uniform around the cell and independent of cell shape

Following Bischofs et al. [18], the radius of curvature of PSF is the result of two competing tensions: R=T/σ, with T the line tension generated in the PSF, and σ the surface tension generated in the cell cortex which pulls perpendicularly on the PSF (Fig. 2a). The cortical tension can thus be expressed as σ =T/R. In other words, surface tension (a global parameter characterizing the 2D traction generated in the cortex) can be, in principle, determined through the measurement of two local scalar parameters characterizing PSF: the radius R, already measured through image analysis, and the line tension T. To measure T, we relied on a technique first introduced in Labouesse et al [28] where a custom-made glass needle, used as a spring of calibrated stiffness, is pushed against the PSF (Fig. 2a). At small PSF indentations, the force applied by the cantilever is equilibrated by geometrical projections of the internal PSF tension T (details in Labouesse et al [28], and SM section 2). Thus T can be retrieved from force-indentation curves of PSF.

**FIG 2.**
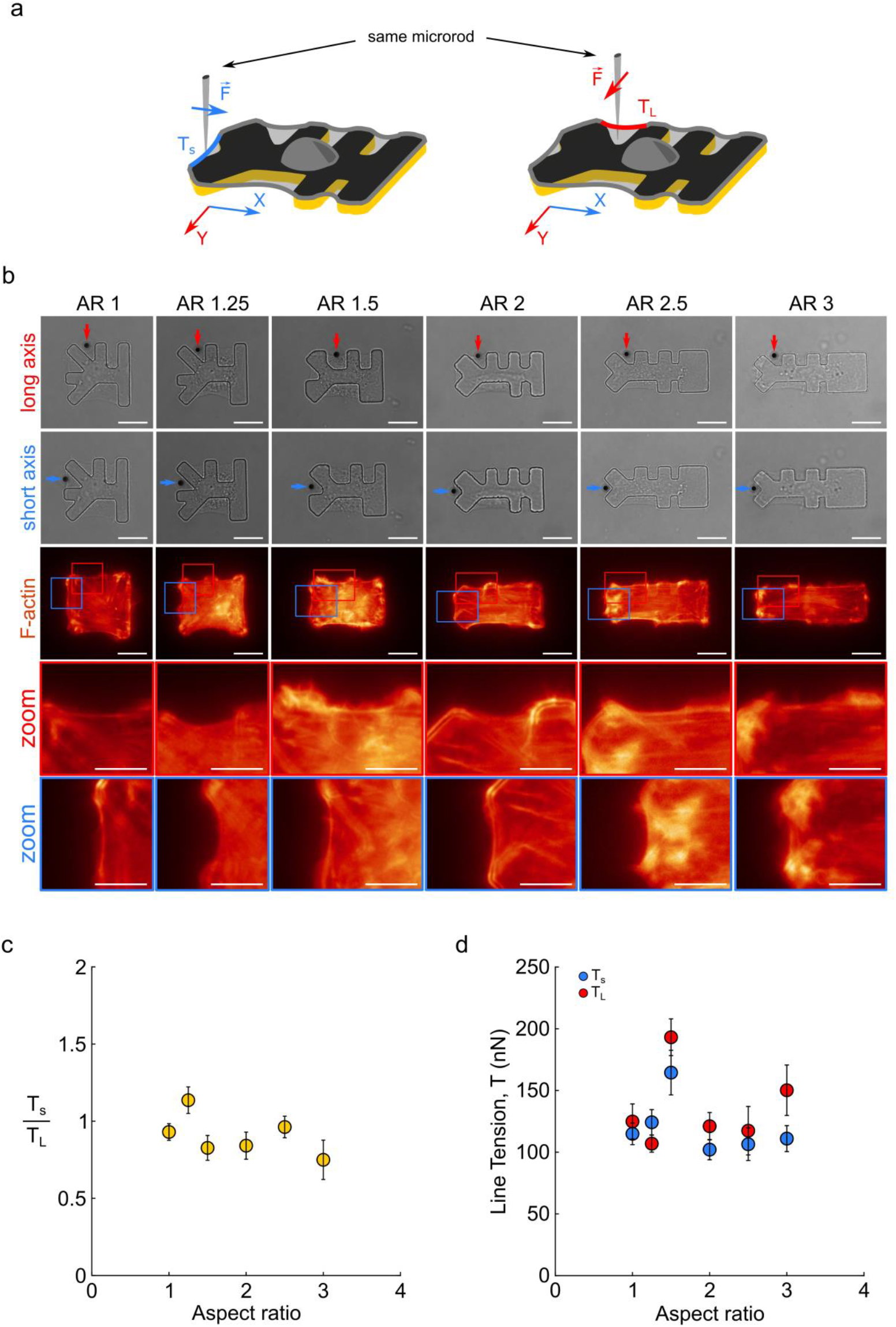
(a) Schematic representation of line tension measurements over peripheral arcs. The same microrod, calibrated along two perpendicular axes, is used to probe the line tension along two peripheral arcs situated along orthogonal cell edges of the same cell. (b) Micrographs (bright-field) of the measurement of the line tension on the long (blue) and short axis (red) for cells spread on micropatterns with aspect ratio ranging from 1 to 3 (scale bar: 20 μm). Blue and red arrows show the microrod positions for the measurements. Micrographs of the actin cytoskeleton (F-actin in red hot), red and blue squares represent the positions of the zoom-in images. Focus on the peripheral arcs of the long (blue) and short (red) axes (scale bar: 10 μm). (c) Values of the line tensions T_s_ (blue) and T_L_ (red) as a functions of the cell aspect ratio. (d) Values of the cell-by-cell ratio 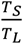 between the line tensions T_s_ (blue) and T_L_ as a function of the cell aspect ratio. (n_AR1_=21, n_AR1.25_=17, n_AR1. 5_=15, n_AR2_=16, n_AR2.5_=13, n_AR3_=15; N_AR1_=2, N_AR1.25_=1, N_AR1. 5_=3, N_AR2_=1, N_AR2.5_=2, N_AR3_=2).

Of note, the cantilever we designed was shaped and calibrated to allow us to measure tension along both the short and long sides of the cells, *i*.*e*. along the x and y axes of the substrate plane (Fig. 2a). As a consequence, we were able to measure, in each individual cell, the tensions T_L_ and T_S_ generated in PSF of the same length but situated along the long and short cell sides respectively (Fig. 2a and 2b). Measuring T_L_ and T_S_ cell-by-cell allows us to make their comparison independent of cell-to-cell variation in tension state (or pre-stress). This can be illustrated on the fluorescence images of Fig. 2b. For instance, one can notice that the cell spread on the micropattern of 1.25 aspect ratio has, along both axes, arc radii that are smaller than those of the cell shown for AR=1. This implies that these cells have very different overall states of tension, independently of anisotropy in tension along the x and y axes.

Thus, thanks to our biaxially calibrated probe, we measured T_s_ and T_L_ for each individual cell. The short axis PSF tension T_s_ and the long axis one T_L_ were found to be approximately equal and to have no noticeable dependence on AR (Fig. 2c and 2d).

### C. Cells with anisotropic shape display anisotropic cortical tension

Having demonstrated that PSF tension T is a constant, and since σ =T/R, the evolution of the cortical tension σ with the cell AR will simply be the inverse of the evolution of R. As a consequence, T_L_ and R_L_ being independent of AR (Fig. 1d and 2c) implies a constant cortical tension σ_y_ along the short axis of the cells for all AR, while constant T_s_ and decreasing R_s_ with increasing AR (Fig. 1d and 2c) indicate an increasing tension σ_x_ along the long cell axis (Fig. 3a, b). Indeed, combining T_s_ and T_L_ measurements with those of R_s_ and R_L_ for each cell, we could determine the ratio σ_x_/σ_y_=[(T_s_ /T_L_)/( R_s_/R_L_)], which represents a cell-by-cell shape-induced tension anisotropy index that should ideally be independent of the overall level of tension or pre-stress in a given cell. This is a key feature of our measurement protocol. Indeed, the overall tension may vary notably from cell to cell as mentioned in the previous section. Thus comparing the mean values <σ_x_=T_s_/R_s_> and <σ_y_=T_L_/R_L_> over cell populations may obscure observation of any significant anisotropy in tension between the x and y axes (see details in SM, section 3).

**FIG 3.**
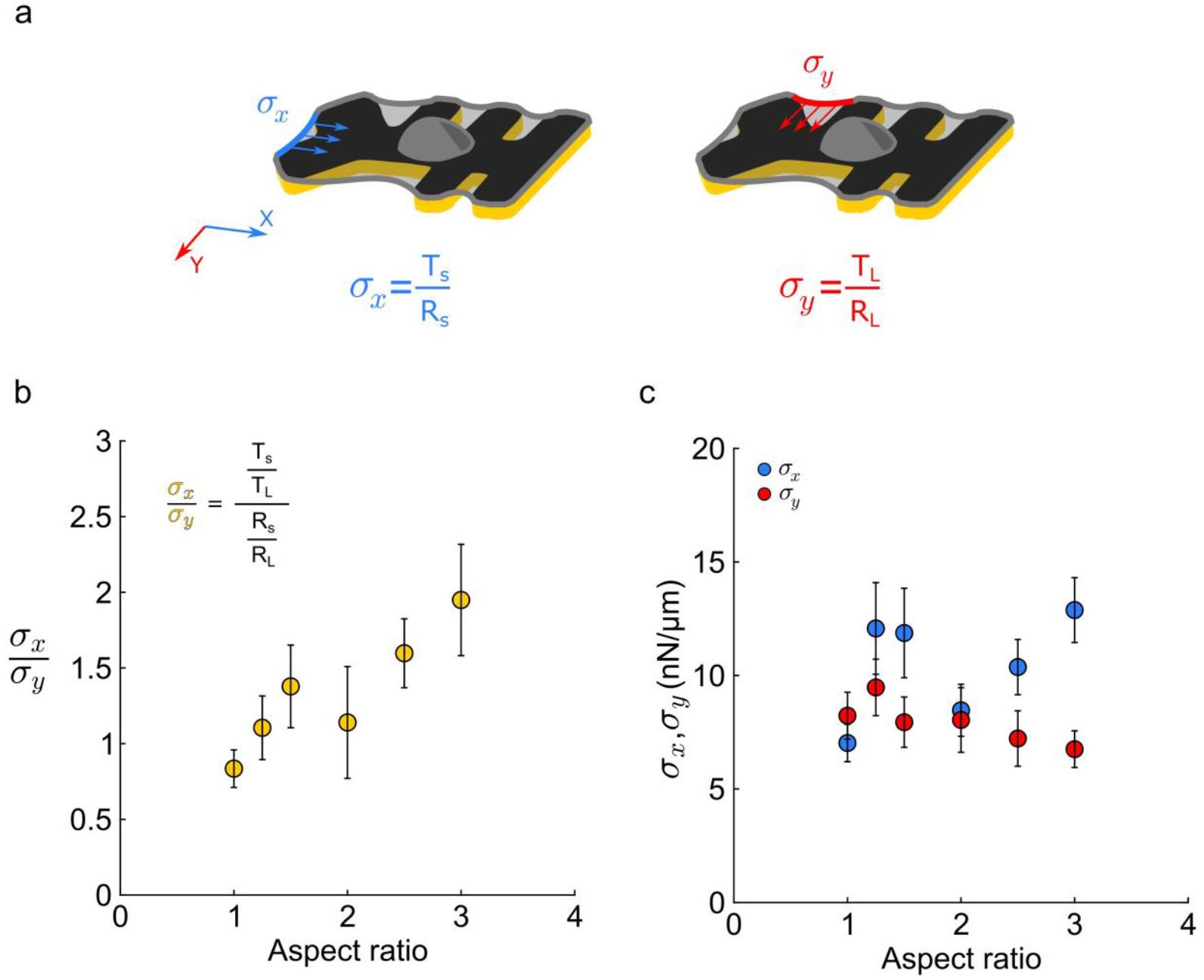
(a) Schematic representation of the cortical (surface) tensions *σ*_*x*_ (blue) and *σ*_*y*_ (red) acting on orthogonal sides of the cells. Surface tensions are calculated as the ratio between the line tensions T and radii of curvature R measured for PSF generated along the specific side the cortical tension is acting on. (b) Values of the surface tensions *σ*_*x*_ (blue) and *σ*_*y*_ (red) as a function of the cell aspect ratio. (c) Values of the cell-by-cell ratio σ_x_/σ_y_ between the cortical tensions as a function of the cell aspect ratio. (n_AR1_=21, n_AR1.25_=17, n_AR1. 5_=15, n_AR2_=16, n_AR2.5_=13, n_AR3_=15; N_AR1_=2, N_AR1.25_=1, N_AR1. 5_=3, N_AR2_=1, N_AR2.5_=2, N_AR3_=2).

Reporting the mean values σ_x_/σ_y_ measured for different cell shape aspect ratios, we could show that σ_x_/σ_y_ increases with cell aspect ratio (Fig. 3b), from a value of ∼1 for AR=1 (i.e. σ_x_ ∼ σ_y_ for “square” cells) to ∼2 for AR=3 (i.e. σ_x_ ∼ 2 σ_y_ for cells 3 times longer than wide). This dependence of the ratio σ_x_/σ_y_ suggests that cell shape anisotropy induces anisotropy in cortical tension, the cortex being more tensed along the longer cell axis. This can also be illustrated by the dependence of the absolute values of the tensions σ_x_ and σ_y_ with the AR (Fig. 3c). Even though these data are more sensitive to cell-to-cell variability as mentioned above. It appears that σ_x_ is globally higher than σ_y_, and that a higher tension along the longer axis is clearly established for the highest aspect ratios.

In sum, using image analysis and non-destructive force measurement combined with modelling, we predict that there exists a shape-induced anisotropy in cortical tension reported in Fig. 3b, c. Thus we reasoned that testing this stress anisotropy with more direct (albeit destructive) measurements of cortical tension, namely through laser ablation, would allow us to validate both this result and the modelling behind it.

### D. Laser-ablation experiments confirm cortical tension anisotropy and its relationship with cell-shape aspect ratio

In order to directly estimate the anisotropy in cortical tension, we used a pulsed infrared laser to generate discoid holes of 5 μm diameter in the cell cortex of cells spread on 2D micropatterns with the same design as those used for line tension measurements. Consistently with an internal tension of the cortex, the initial circular hole expanded and reached a steady shape after typically 10 to 20 seconds (Fig. 4a). We found that the final shape was increasingly elliptic for larger cell aspect ratio, and that the major axis of the elliptic holes was increasingly oriented towards the longest cell axis (Figure 4b). Qualitatively, these observations suggest that tension is anisotropic in the cell cortex, cortical tension being maximum (noted σ_max_) along the major ellipse axis, and minimum (σ_min_) along the minor axis (Fig. 4b). The open holes being more and more elliptic when the aspect ratio is increased suggests that cortical anisotropy is increasing with the anisotropy of the cell shape.

**FIG 4.**
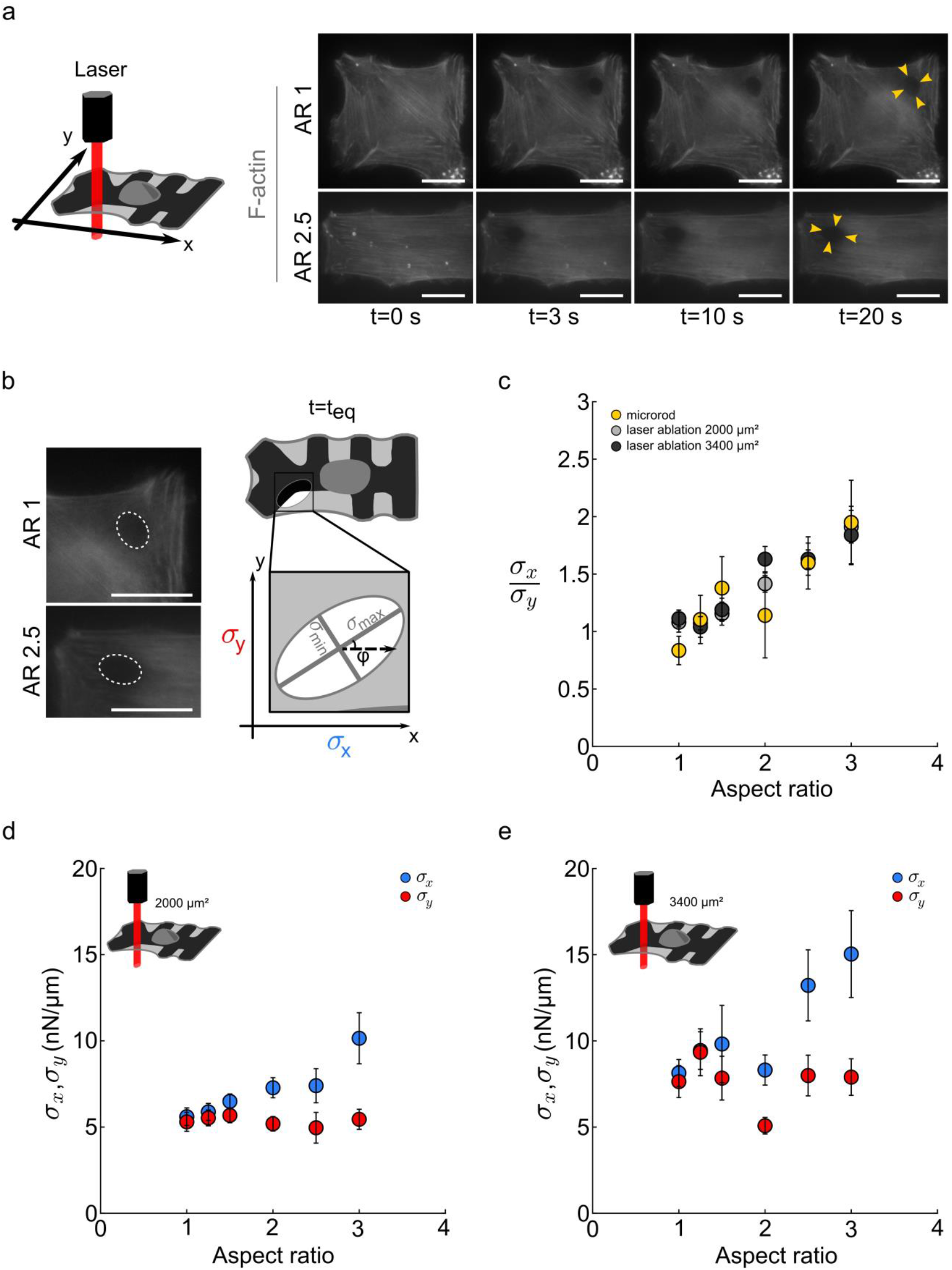
(a) Laser ablation experiment. On the left, schematic representation of a laser ablation. Cells are spread on 2D micropatterns. A 5 μm diameter hole is performed with a pulsed infrared laser far enough from the nucleus and the edges of the cells. On the right, typical laser ablation experiments done on aspect ratios 1 and 2.5. After around 15 s, the initial hole relaxes into a steady ellipse. Scale bars: 20 μm. (b) On the left, zoom of the relaxed ellipses (Scale bars: 20 μm). On the right, sketch of the method used to extract surface tension from laser ablation. Major and minor axis of the relaxed ellipse are proportional to the two main stresses in the probed material (see SM section 3). These main stresses can be projected along the x and y axis (see Appendix B). (c) Mean values of the ratio of the surface tensions 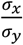 as a function of the aspect ratio for both laser ablation (light and dark grey dots, corresponding to spread areas of 2000 and 3400 μm^2^, respectively) and microrod (yellow dots) experiments. (d) Mean values of surface tensions σ_x_ and σ_y_ as a function of aspect ratio for cells spread on micropatterns of 2000 μm^2^ (d) and 3400 μm^2^ (e). (For micropatterns of 2000 μm^2^: n_AR1_=19, n_AR1.25_=20, n_AR1. 5_=22, n_AR2_=34, n_AR2.5_=22, n_AR3_=15; N=2 for AR=1, N=1 otherwise. For micropatterns of 3400 μm^2^: n_AR1_=23, n_AR1.25_=12, n_AR1. 5_=9, n_AR2_=26, n_AR2.5_=21, n_AR3_=13; N N=1).

In order to quantify these observations, we adapted the model developed in [29,30] and considered the cortex as a material of mean thickness *e* and Young’s modulus *E* displaying an anisotropic in-plane internal stress varying between a minimum value *τ*_*min*_ and a maximum value *τ*_*max*_. It is then possible to express *τ*_*min*_ and *τ*_*max*_ as functions of *E* and the measured lengths *a* and *b* of the major and minor axes of the elliptic holes (Appendix A). The 2D maximal and minimal cortical tensions *σ*_*max*_ and *σ*_*min*_ can then be expressed as *σ*_*max*_ = *eτ*_*max*_ and *σ*_*min*_ = *eτ*_*min*_. Since the cortical effective Young modulus of spread 3T3 cells was measured at 20 kPa [31] and cortical thickness reported to be in the range 350-600 nm [32,33], we took *e*=0.5 μm and *E*=20 kPa to estimate σ_min_ and σ_max_ for each cell submitted to laser ablation. Finally, these σ_min_ and σ_max_ values were projected along the long and short axes of the cells to determine the values σ_x_ and σ_y_ of the cortical tensions acting respectively along these axes (Appendix B). Thus we could get, from laser ablation, an independent estimation of σ_x_/σ_y_ for each tested cell. We found that σ_x_/σ_y_ increased with AR, with values in very good agreement with those retrieved from line tension and curvature radii measurements (Fig. 4c). This agreement between cortical tension ratios obtained from independent measurements and interpreted through different physical models, also stands in favour of the dependence of the absolute values of the cortical tensions σ_x_ and σ_y_ on the cell aspect ratio (Fig. 4d and Fig. 3c), confirming that σ_y_ is constant, while σ_x_ is an increasing function of the cell aspect ratio. Remarkably, the absolute values σ_x_ and σ_y_ retrieved from laser ablation are of the same order of magnitude as those measured with the microrod technique.

Of note, the ratio σ_x_/σ_y_ is independent from the spread area of the cells (Fig. 4c). Consistently, on micropattern of larger area cells display the same evolutions of σ_x_ and σ_y_ with cell AR, but the absolute values of σ_x_ and σ_y_ are shifted toward higher values (Fig. 4e).

### E. Analysing the relationship between cortical SF (CSF) orientation and tension anisotropy reveals an increasing contribution of CSF to cortical tension with increasing cell aspect ratio

In the previous sections, we showed, by two independent measurement techniques and analyses, that cortical tension becomes increasingly anisotropic as the cell aspect ratio is increased. We then asked how cell AR could lead to anisotropy in cortical tension. To answer this question, we focused on the structure of the actin cortex, searching for a possible structural anisotropy controlled by the cell aspect ratio.

Cortical tension is known to be mainly due to actomyosin-based contraction [8,15,34,35]. Anisotropic actomyosin tension is generally associated with the anisotropy of the actomyosin meshwork itself [3,36]. Thus, we examined the orientation of cortical stress fibres (CSFs). CSFs act as force dipoles, embedded in an isotropic cortical mesh [16], leading to spatially concentrated and oriented forces. Since CSF orientation has been reported to depend on cell shape [14,16,19,20,24], we hypothesized that such an AR-dependent orientation of CSF could lead to the observed cortical tension anisotropy.

In order to test this hypothesis, we first analysed the orientation of actin filaments in the context of both experimental protocols, *i*.*e*. microrod (MR) and laser ablation (LA) measurements (Figure S1a). We found that CSFs increasingly orient along the long axis as the AR increases (Fig. S1a-b), thus being correlated with the increase of cortical tension anisotropy (Figure S1c). In the case of laser ablation, comparing the local polar distribution of the actin filaments *before ablation* with the polar distribution of the major axis of the elliptical holes obtained *after ablation*, these distributions appear visually the same (Figure 5a-b), an observation also valid for the larger cell area of 3400μm^2^ (Figure S2). The major axis of the ellipses corresponding to the direction of the maximal tension; it appears that this direction corresponds to the most probable orientation of the CSF, suggesting a direct relationship between CSF orientation and tension anisotropy.

**FIG 5.**
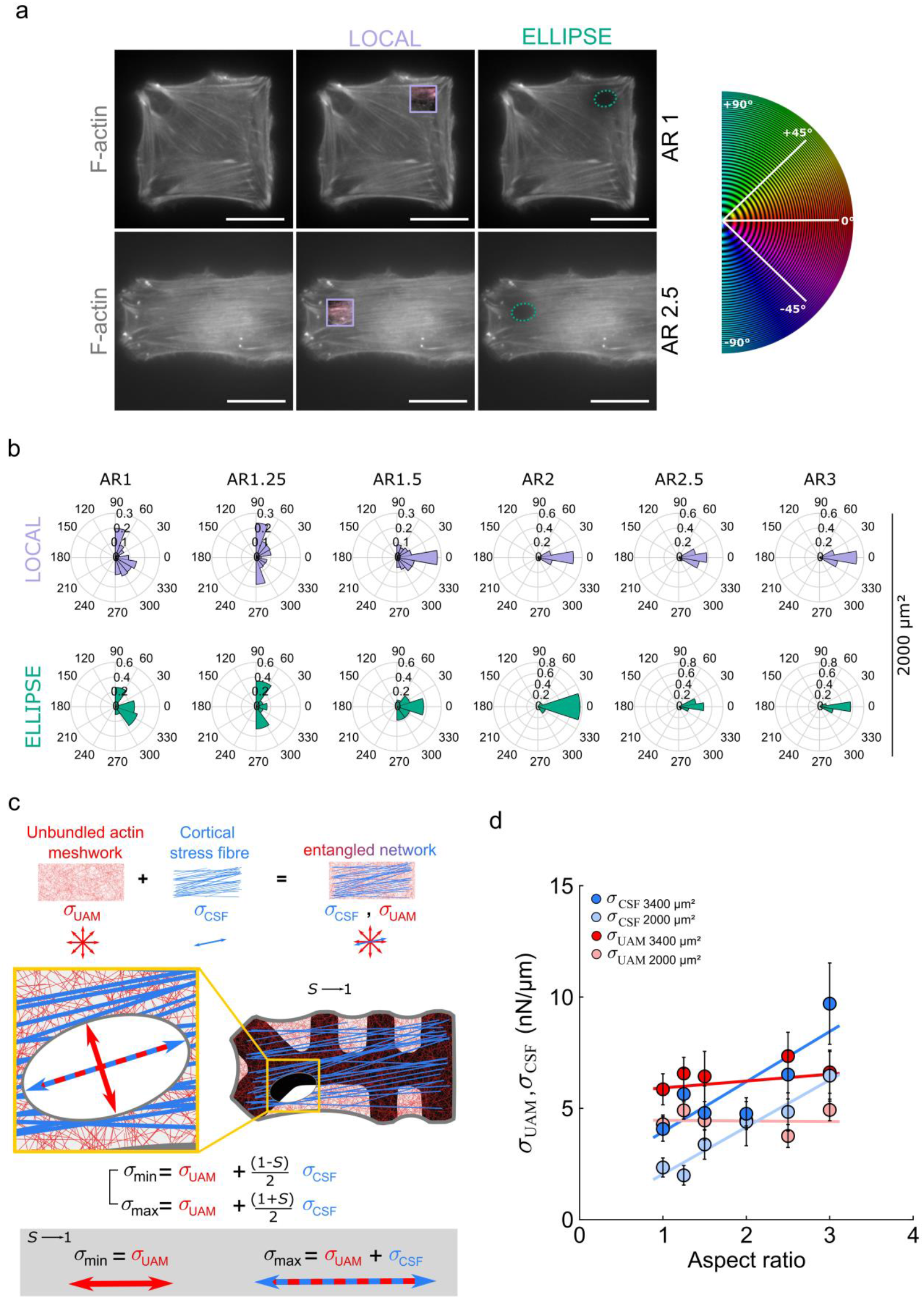
(a) Micrographs of actin cytoskeleton (F-actin, grey) for cells spread on micropatterns of anisotropy 1 and 2.5. In order to determine the local orientation of CSF, we cropped a 10 μm square centred on the discoidal ablated region (in purple) and measured stress fibres orientation with OrientationJ in this region. For ellipse orientation, we calculated the angle between the major axis *b* of the ellipse and the x-axis of the micropatterns. (b) Angle distribution of local stress fibres (purple) and ellipse openings (green) for 2000 μm^2^ micropatterns(c) Schematic illustrating the relationships between the main surface tensions σ_min_ et σ_max_ obtained from the opening of the ellipses, and the cortical tensions σ_UAM_ et σ_CSF_. (d) Evolution of the contribution to surface tension of the isotropic cortex σ_UAM_ (in red) and of the cortical stress fibres σ_CSF_ (in blue) as a function of cellsaspect ratio. Light and dark colours correspond to 2000 and 3400 μm^2^ micropatterns, respectively.

In order to make this relationship quantitative, we relied on a 2D physical model of the cortex. The cortical meshwork is described as a composite material with an isotropic mesh spanning the cell surface, which we will call the *unbundled actin meshwork* (UAM), embedding a network of denser and oriented actin structures, presumably bundled, namely the CSF [16,23,37,38]. We then decompose the stress tensor in the cortex as **σ = σ**_uam_ + **σ**_csf_, that is, the sum of the stress in the UAM and in the CSF. The UAM being isotropic, we write **σ**_uam_ = σ_uam_ **I**, thus corresponding to a (negative) surface pressure, with the scalar σ_uam_ expressed in Pa·m.

The situation is quite different for the stress generated by the CSF. Individual CSF act as active force dipoles, the force being applied along the CSF axis, and thus their contribution to tension is oriented. However, the overall stress generated by a collection of CSF can be either isotropic or anisotropic depending on the relative orientations of the CSF. For instance, if the CSF orientations are randomly distributed, then the stress due to CSF activity will be isotropic (nematic order parameter magnitude *S=0*). Conversely, if all CSF are oriented along the same direction (*S=1*), the stress will be anisotropic and oriented along CSF direction. The general case corresponds to an intermediate situation where CSF are partially oriented (*0<S<1*). Then, the stress arising from the CSF contains isotropic and anisotropic components (Appendix C). As shown in the Appendix, the anisotropic part of this CSF-generated stress is proportional to *S*. The minimal and maximal stresses generated in the composite cortex (*i*.*e*. UAM + CSF) are then given by

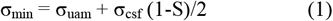

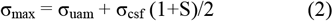

Note here two interesting limits. First, when CSF are randomly oriented, *S*∼0 and σ_min_ ∼ σ_max_ ∼ σ_uam_ + σ_csf_ /2. This is the case for the lowest AR with laser ablation leading to almost circular holes and <σ_x_/σ_y_> ∼ 1 (Fig. 4c). Second, when all the fibres are oriented along the same direction, *S* ∼ 1, leading to σ_min_ ∼ σ_uam_ and σ_max_ ∼ σ_uam_ + σ_csf_ (scheme in Fig. 5c). Indeed, for the highest AR, the CSF are oriented along the long axis of the cells (the x-axis) and we found <σ_x_/σ_y_> ∼ 2 (Fig. 4c), suggesting that σ_csf_ ∼σ_uam_ in that case. By inverting Eqs. (1) and (2) one can express σ_uam_ and σ_csf_ as functions of σ_min_, σ_max_, and *S*:

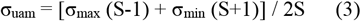

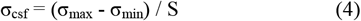

Thus, from the measurements of σ_min_, σ_max_, and S for each AR value, we are able to determine the values of the specific contributions of the isotropic mesh (σ_uam_) and that of the stress fibres (σ_csf_) to the overall cortical tension as function of the cells AR (Fig. 5d). We find that σ_uam_ is almost independent of the cell AR, while σ_csf_ increases with the AR. Of note, this trend is the same independently of the overall cell spread area, but the absolute values of σ_uam_ and σ_csf_ are shifted toward higher values when the spread area is increased (Fig. 5d).

## IV. CONCLUSIONS AND DISCUSSION

By designing substrates with original 3D micro-patterns, we reveal the effect of the 2D aspect ratio (AR) of spread single cells on the structure and properties of their actin cortex. These micro-patterns allowed us to vary the AR while maintaining constant cell area, free perimeter and the length of peripheral stress fibres (PSF). Combining these micro-patterns with *in situ* measurements of the tension of single PSF, we first show that an anisotropic cell shape leads to anisotropy in cortical tension, the longest axis of the cell exhibiting higher tension, and tension anisotropy increasing with cell AR. This shape-driven anisotropy in cortical tension was independently confirmed by laser ablation experiments. Moreover, by measuring the orientations of cortical stress fibres (CSF) for different cell AR, we were able to make explicit the relationship between CSF organisation (mean orientation and order parameter) and the anisotropy of cortical tension. In particular, describing the cortex as made of CSF (force dipoles) embedded in an isotropic contractile cortex, which we name the unbundled cortical meshwork (UAM), we could estimate the specific contributions of the isotropic mesh (σ_uam_) and that of the stress fibres (σ_csf_) to the overall cortical tension as function of the cells AR.

Interestingly, for the highest AR tested, CSF were almost parallel to the longest axis of the cell and we found that the contributions of the UAM and that of the CSF were of equivalent magnitude (σ_csf_ ∼ σ_uam_). This finding can be compared to the observation done by Vignaud et al. on cells plated on dumbbell-shaped micro-patterns with AR = 5 [16]. In this configuration, cells exhibited two main peripheral stress fibres and only very few small and thin fibres otherwise. By combining photoablation with traction force microscopy, the authors could show that each of the main stress fibres contributed to 25% of the traction energy, e.g. the two main stress fibres contributed to 50% of the traction energy, the remaining 50% being thus due to the fine actin meshwork, in line with our own finding of σ_csf_ ∼σ_uam_.

Of note, the cell-shape-based anisotropy of cortical tension we measured here for the first time is reminiscent of the mechano-sensitivity of actomysosin itself [39–41]as previously suggested in the context of cell spreading and polarization on substrates of anisotropic stiffness [42]. It also poses the question of how CSF form [43]. In particular, it has recently been shown that stochastic contractile pulses induce the formation of cortical stress fibres in spread cells [44]. This kind of process might be at play during shape establishment when cells spread on substrates of anisotropic shapes, and where oriented transient increase in cortical tension could generate accordingly oriented stress fibers [45,46].

As a conclusion, we have demonstrated here that cell shape by itself can control the material properties of the cell cortex, leading to specific sub-cellular organization and anisotropy in tension. This shape-induced modulation of the cell structure and contractile activity could be of importance in fundamental biological processes such as development where a shape-based oriented reinforcement and anisotropy of tension could feedback on the genetic programs [47], acting thus as a quality control to ensure a proper morphogenesis [1,3,48]

## ACKNOWLEDGEMENTS

The study was supported by the labex “Who AM I?”, labex ANR-11-LABX-0071, as well as the Université Paris Cité, Idex ANR-18-IDEX-0001, funded by the French Government through its “Investments for the Future” program.

## AUTHOR CONTRIBUTIONS

F.F. designed and performed all experiments, analysed and interpreted the results, and wrote the paper; S.A. supervised the experimental work, helped with data analysis, interpreted the results, and wrote the paper; A.P. carried out preliminary 3D-micropattern experiments; C.D.P. gave technical support with laser ablation experiments; A.R. helped for cell culture; C.V. helped with plasmid production for LifeAct 3T3 cell line; J.F., helped with data interpretation and discussions; H.E., provided critical feedback; J.E., A.C.-J., helped with the tensorial formalism; M.T. provided ideas, assisted data interpretation and discussions; D.P. designed the experiments, helped with experimental techniques, with data analysis and interpretation of the results, and wrote the paper; A.A. conceived and supervised the study, designed the experiments and modelling, analysed and interpreted the results, wrote the paper, and sought fundings. All the authors reviewed, edited and approved the paper.

## APPENDIX A: SURFACE TENSION EXTRACTION FROM LASER ABLATION EXPERIMENTS

To extract the surface tension from laser ablation experiments, we used the formalism developed for hole drilling in materials with anisotropic pre-stress [29,30]. We performed a 5 μm diameter discoidal ablation (hole) in the cell cortex. At equilibrium, we measure the major and minor axis of the ellipse, *a* and *b*, respectively (fig. 6). We retrieve the stresses *τ*_*max*_ and *τ*_*min*_ from the displacements, along any point *M* of the ellipse and oriented along the vector ***e***_*r*_, following the relation:

**FIG 6.**
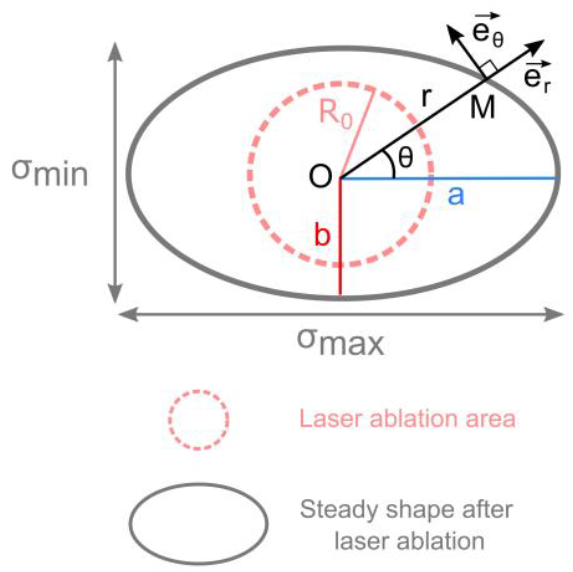
Schematic representation of the laser ablation/hole drilling method. At equilibrium, the initial hole (pink dashed line) becomes an ellipse (grey line) with major and minor axis a (blue) and b (red), respectively.

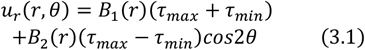

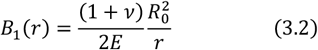

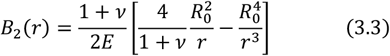

Where *v* is the Poisson ratio of the cell cortex, *E* its Young Modulus, *R*_0_ the initial radius of the photoablated area, and *r* the length of the vector ***OM*** (see fig. 6). *τ*_*max*_ and *τ*_*min*_ are calculated from the displacements *u*_*max*_ = *a* − *R*_0_ and *u*_*min*_ = *b* − *R*_0_ that can also be expressed from equation (3.1) for *θ* = 0 and 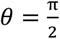, respectively (see figure 6), this gives:

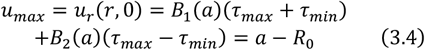

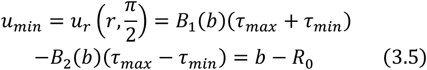

Then,

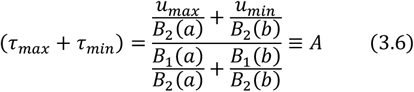

And

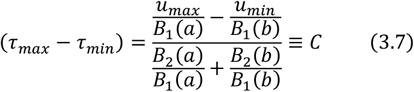

Finally,

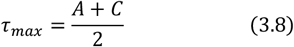

And

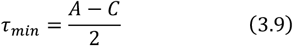

Also, expanding *A* and *C* to show their dependence on the geometry of the hole (*R*_*0*_, *a, b*) and the mechanical features of the cortex (*E*,υ), one finds:

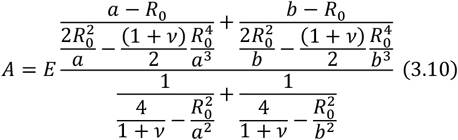

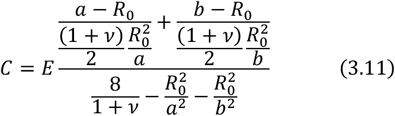

Finally, surface tensions can be expressed as follows:

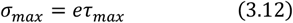

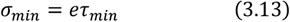

Where *e* is the thickness of the cortex. We took *e* = 0.5 μ*m*, in agreement with data from the literature (Kumar et al. 2019; Smeets et al. 2019).

## APPENDIX B: DETERMINATION OF THE CORTICAL TENSIONS *σ*_*x*_ AND *σ*_*y*_ FROM *σ*_*max*_ AND *σ*_*min*_

After laser photoablation, the major and minor axes of the ellipses are proportional to the two main stresses, *τ*_*max*_ and *τ*_*min*_, as well as to the two cortical tensions in the cell cortex, *σ*_*max*_ and *σ*_*min*_, respectively (eqn. 3.12 & 3.13). Since an angle ϕ between the major axis of the ellipse and the long axis x (figure 7). Those tensions can be modelled as stress tensors which can be projected onto the long axis x and the short axis y of the cell as follows:

**FIG 7.**
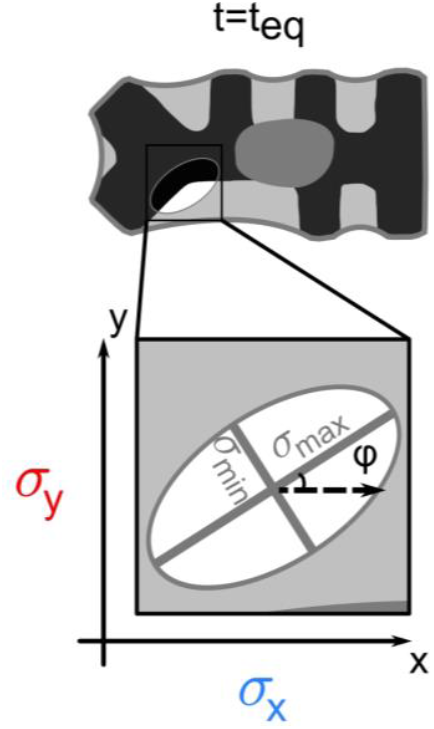
Schematic view of the cell cortex, at equilibrium, after laser photoablation. The major and minor axis are oriented along the two main tensions σ_max_ and *σ*_*min*_. φ is the angle between *σ*_*max*_ and the long axis *x*.

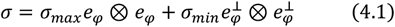

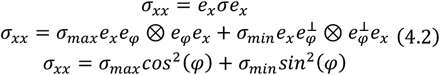

And

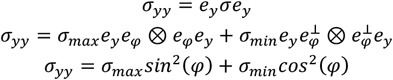

## APPENDIX C: EXPLICIT MODEL FOR A TWO COMPONENT CORTICAL TENSION

The cortex is described as a composite material with an isotropic mesh spanning all the cell surface (that we will from now on call the cortex) displaying in some locations a denser and oriented actin structures, e.g. the CSF [16]. As a surface tension, the cortical tension can be described as a 2D negative pressure, a force per unit length. The cortex, e.g. the isotropic mesh, leads to an isotropic tension *σ*_*cort*_: if one makes a cut along a line of length *l* in the cortex, whatever its orientation, the force applied on this line will be simply *σ*_*cort*_ *l*. The situation is quite different when it comes to the forces applied by the CSF. The CSF act as force dipoles, the force being applied along the CSF axis, and their contribution to tension is also oriented. In that case, the forces applied to an imaginary line-cut of length *l* depends on the relative orientations of the line and the CSF, the force generated by each fibre being projected on the normal to the line (Fig. 8c).

**FIG 8.**
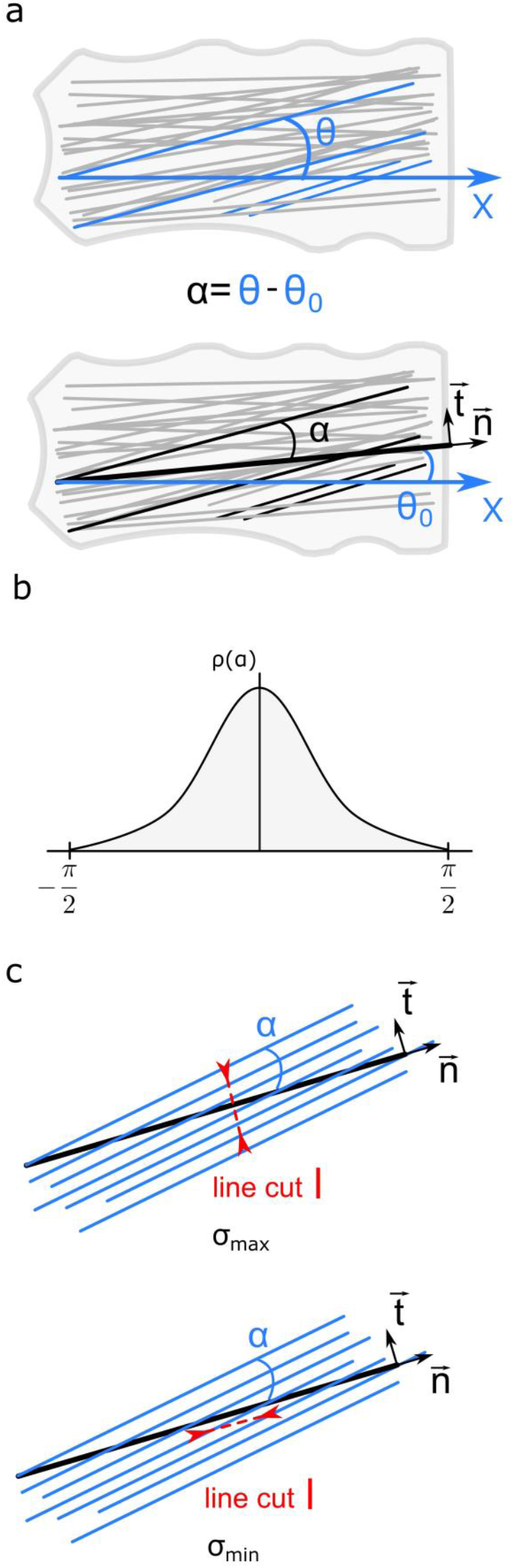
(a) Schematic representation of the angles *θ, θ*_0_ and *α*. *θ* is the angle between the axis *x* of the micropattern and the axis of a given CSF (or a group of CSF of the same orientation comprised between *θ and θ* + *dθ*). *θ*_0_ is the angle between the axis *x* of the micropattern the axis of the nematic director ***n***, and *α* the one between a given CSF and the nematic director ***n***. (b) Schematic representation of the angular density/distribution function of the CSF. (c) Schematic representation of the two line cuts of length *l*, one parallel to ***t*** (to compute *σ*_*max*_) the other parallel to ***n*** (for *σ*_*min*_ determination)

For instance, if the cut is normal to a given fibre, then all the force will be acting to open the cortex; while a cut made parallel to a fibre will obviously not be submitted to any force and will remain closed.

We note *θ* the angle between the axis of a given CSF and the *x* axis of the micropatterns, e.g. le longest axis for *AR* > 1, and *θ*_0_ the angle of the nematic director, e.g. the mean value of *θ* over all CSF: *θ*_0_ =< *θ* >. Let’s note *α* the angle between the axis of a given CSF and the nematic director, *α* = *θ* − *θ*_0_. By definition *α* values are distributed around 0 (fig. 8a-b). Let us consider a subset of the CSF the orientation of which is comprised between *α* and *α* + *dα*. To determine the contributions of this subset of CSF to the cortical tensions *σ*_*max*_ and *σ*_*min*_ applied along (vector ***n***) and orthogonally (vector ***t***) to the nematic director respectively, we consider two line cuts of length *l*, one parallel to ***t*** (to compute *σ*_*max*_) the other parallel to ***n*** (for *σ*_*min*_ determination), fig. 8c.

### 1. Contribution of the CSF to σ_max_

If we note N the total number of CSF and 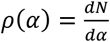 the subset of the CSF the orientation of which is comprised between *α* and *α* + *dα*, then N is given by:

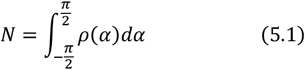

The line density of the subset of CSF oriented along the direction *α*, is given by 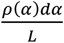 with *L* a typical length. Then, the number of CSF pulling on a line cut of length *l* oriented orthogonally to ***n*** is given by:

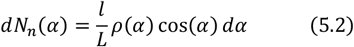

If we note ***f*** the force exerted by one CSF, then the force exerted on the line cut (e. g. its projection along ***n***) is given by:

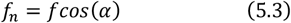

Thus, the force *dF*_*n*_(*α*)exerted by the subset of CSF oriented along *α* is given by:

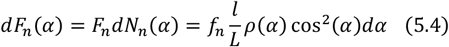

and the corresponding contribution to the maximum cortical tension *σ*_*max*_(along the nematic director) then writes:

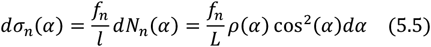

with 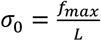 the typical surface tension produced by a single fibre, leading to:

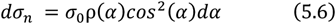

Thus, the contribution of the subset of CSF to the maximal surface tension, e.g. along the director ***n***, is given by:

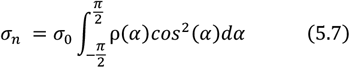

e.g.

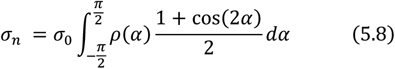

leading to:

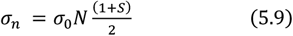

with *σ*_*csf*_ = *σ*_0_*N*, the surface tension due to all the CSF and *S* the nematic order parameter.

Finally, the maximum surface tension *σ*_*max*_, is the sum of the contributions of the isotropic meshwork *σ*_*cort*_ and that of the CSF *σ*_*n*_, e.g.

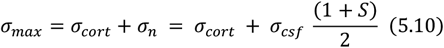

### 2. Contribution of the CSF to σ_min_

The contribution of the CSF to *σ*_*min*_ due to the forces acting on a line cut parallel to ***n***, e.g. the projections along ***t*** of the forces generated by the CSF. Equations (5.2) to (5.7) must then be rewritten by replacing *cos*(*α*)by *sin*(*α*), leading to:

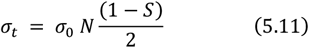

And finally,

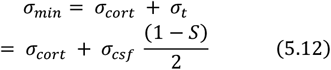

### 3. Comparison with the usual tensorial formalism

While we decided to explicitly write the contributions of the actin meshwork *σ*_*cort*_ and that of the stress fibres *σ*_*csf*_ to express the minimal and maximal principal tensions *σ*_*min*_ and *σ*_max_ (equations (5.10) and (5.12), corresponding to the minor and major axes of the ellipses opened after laser ablation respectively), their expressions are in total agreement with the expressions derived usually from the tensorial description of nematic liquid crystals [14,49]. For instance, in Schakenraad et al., *σ*_*min*_ and *σ*_*max*_ are written as:

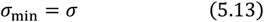

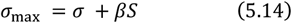

where *σ* accounts for all isotropic contributions to cortical tension, *S* the order parameter and *β* a coefficient having the dimensions of a surface tension. Equations (5.10)-(5.12) are equivalent to (5.13)-(5.14) provided that *β* = *σ*_*csf*_ and 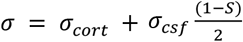 which indeed represents all the isotropic contributions to surface tension, e.g. the one from the isotropic actin meshwork *σ*_*cort*_, plus the isotropic contribution of the stress fibres 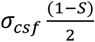; this last contribution being simply 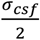 when the stress fibres are randomly oriented (*S* = 0), and null when all fibres are oriented along the same direction (*S* = 1).

## Notes

### Competing Interest Statement

The authors have declared no competing interest.

### Summary of Updates

Title of appendix B has been modified.

