## Supplementary Material for "Disentangling the contributions of stress fibres and the unbundled actin meshwork to the anisotropy of cortical tension in response to cell shape"

#### 1. Effect of free length on PSF radius, correlation between opposite sides

Of note, the specific design of our micropatterns also allowed us to test the effect of the free length between adhesive regions on the curvature of PSF. Indeed, in addition to the non-adhesive regions of length $d_1$=15 µm mentioned above, all micropatterns, independently from their AR, were designed with additionally a non-adhesive region of length $d_2$=30 µm, also situated along the long axis of the cell, but situated on the opposite side of the pattern (Fig. 1). In other words, we could compare the radii of PSF generated along the long axis of the same cell, but for free lengths of different values (Fig. 1). In these conditions, for all aspect ratios, the radius of curvature of PSF is larger for the larger free length (Fig 1d, e). On average, we found $\langle R_{L15} \rangle$ = 20 µm for $d_1$=15 µm, and $\langle R_{L30} \rangle$=50 µm for $d_2$=30 µm. This behaviour is in line with the observations reported in Bischofs et al[1]. Consistently with the observation made with the short free length of 15 µm, increasing the cell AR has no effect on the curvature radius of PSF formed on 30 µm free length. Remarkably, in line with the hypothesis that cortical tension could be the parameter allowing le cell AR (e.g. a global cell-scale parameter) to control local features such as the PSF curvature radius, we noticed that the fluctuations observed on the values of $\langle R_{L15} \rangle$ and $\langle R_{L30} \rangle$ for different AR values where quite similar (Fig. 1e) suggesting a link at distance between opposite sides of the cells. We suggest that this can be a signature of cortical tension fluctuations from sample to sample revealed by PSF curvature variations on opposite sides of the cells.

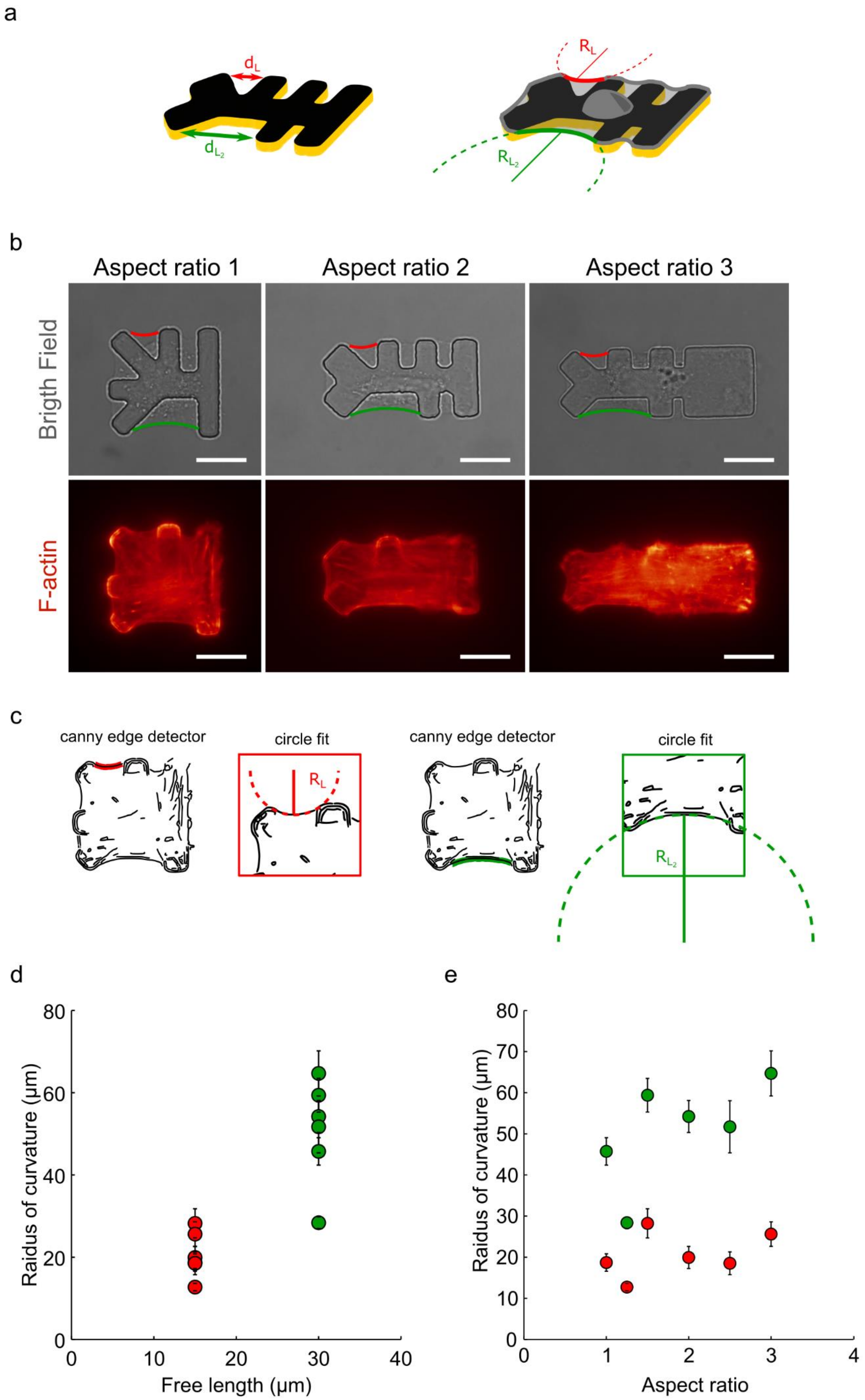
a
d_L
d_L2
R_L
R_L2
b
Aspect ratio 1
Aspect ratio 2
Aspect ratio 3
Brigth Field
F-actin
c
canny edge detector
circle fit
R_L
canny edge detector
circle fit
R_L2
d
Raidus of curvature (μm)
Free length (μm)
0
10
20
30
40
0
20
40
60
80
e
Raidus of curvature (μm)
Aspect ratio
0
1
2
3
4
0
10
20
30
40
50
60
70
80

**Figure 1:** (a) Schematic representation of a 3D micropattern with an aspect ratio of 2 and two different free lengths $d_L$ (red) and $d_{L_2}$ (green), measuring 15 and 30 µm, respectively. (b) Bright field and fluorescence (F-actin, red hot) images of cells spread on micropatterns of 2000 µm² with an aspect ratio ranging from 1 to 3. Free lengths of 15 and 30 µm are represented in red and green, respectively. (c) Procedure of arc radius fitting. First, the cell contours are segmented with a canny edge detector filter. Then, both ends of each stress fibre are manually determined and a circle is fitted to the arc fibre using the Pratt's Method. (d-e) Radius of curvature of the peripheral arcs as function of the free length (d) and of the aspect ratio (e). (number n of cells $n_{AR1}$=21, $n_{AR1.25}$=17, $n_{AR1.5}$=15, $n_{AR2}$=16, $n_{AR2.5}$=13, $n_{AR3}$=15, number N of independent experiments $N_{AR1}$=2, $N_{AR1.25}$=1, $N_{AR1.5}$=3, $N_{AR2}$=1, $N_{AR2.5}$=2, $N_{AR3}$=2).

#### 2. Line tension measurement in a peripheral stress fibre

Cells are spread on micropatterns with fibronectin gaps (fig. 2a-b). These non-adherent areas induce the formation of peripheral actin arcs. In addition, the micropatterns are three-dimensional, e.g. designed to be 5 µm in height. Thus, we end up with elevated isolated actin arcs which we will be able to mechanically probe without any friction between the arcs and the substrate. We push on the stress fiber with a glass cantilever (micro-rod) of known stiffness $k$ with a $1\mu m/s$ ramp for $5s$ (fig. 2c-d). The base $B$ of the cantilever is moved by $5\ \mu m$ in total and we detect the position $p$ of its tip over time. The force with which we push on the stress fibre is then given by:

$$F = k(B - p) = k\delta \quad (1.1)$$

Force balance consideration gives the inner tension of the stress fibre as follows:

$$T = \frac{F}{2\sin(\varphi)} \quad (1.2)$$

Geometrical considerations lead to:

$$sin(\varphi) = (x_0 + e)\sqrt{\frac{1}{d^2 + (x_0 + e)^2} - \left(\frac{1}{2R}\right)^2} - \frac{d}{2R} \quad (1.3)$$

where $R$ is the radius of curvature of the PSF, $e$ the PSF indentation, $d$ the distance between the position of the applied force and the PSF adhesion site and $x_0$ the height of the arc (for more details see [2]). The line tension is then extracted from the linear part of the force curves (fig. 2g-h).

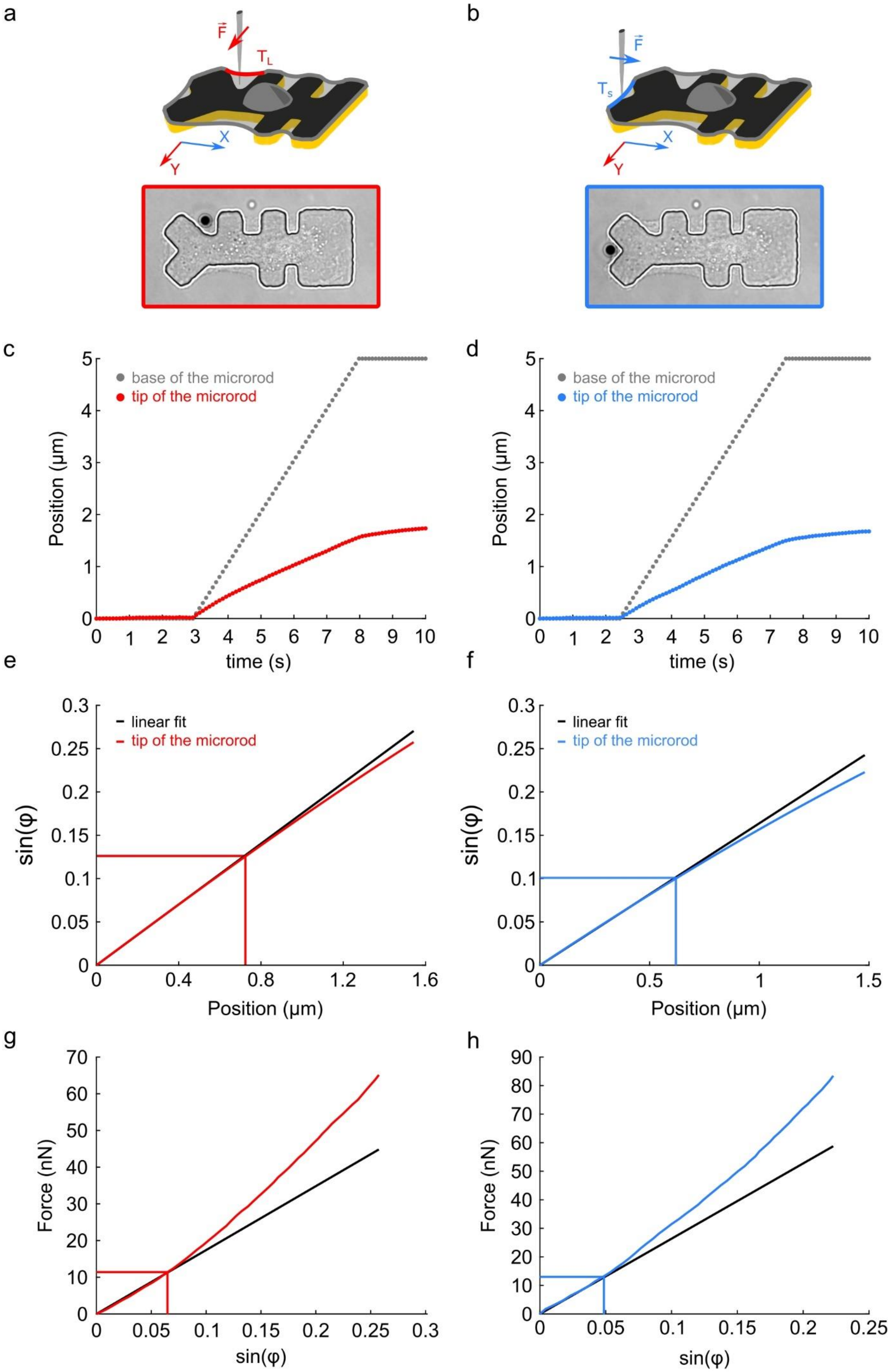
a
b
F
T_L
T_s
X
Y
c
d
base of the microrod
tip of the microrod
Position (µm)
time (s)
e
f
linear fit
tip of the microrod
sin(φ)
Position (µm)
g
h
Force (nN)
sin(φ)

**Figure 2:** Method to extract the stress fibre line tension from the micro-rod experiment. (a-b) Schematic representation of the experimental procedure of line tension measurement of the peripheral stress fibres (PSF). (c-d) Position of the tip $p$ and base $B$ of the micro-rod in time for respectively the longitudinal and lateral stress fibres. (e-f) $sin(\varphi)$ as a function of the position of the microrod tip for respectively the longitudinal and lateral stress fibres. Beyond 0.6 µm indentation, the relationship is not linear anymore, indicating the onset of the elastic regime where the contribution of fibre elastic elongation to tension cannot be neglected. (g-h) The force $F$ applied to the stress fibres as function of $sin(\varphi)$ for respectively the longitudinal and lateral stress fibres. The slope of the linear part is equal to $2T$, where $T$ is the pre-existing active tensions generated by actomyosin in the stress fibres.

#### 3. Line tension ratios: cell by cell measurements vs estimation over a population

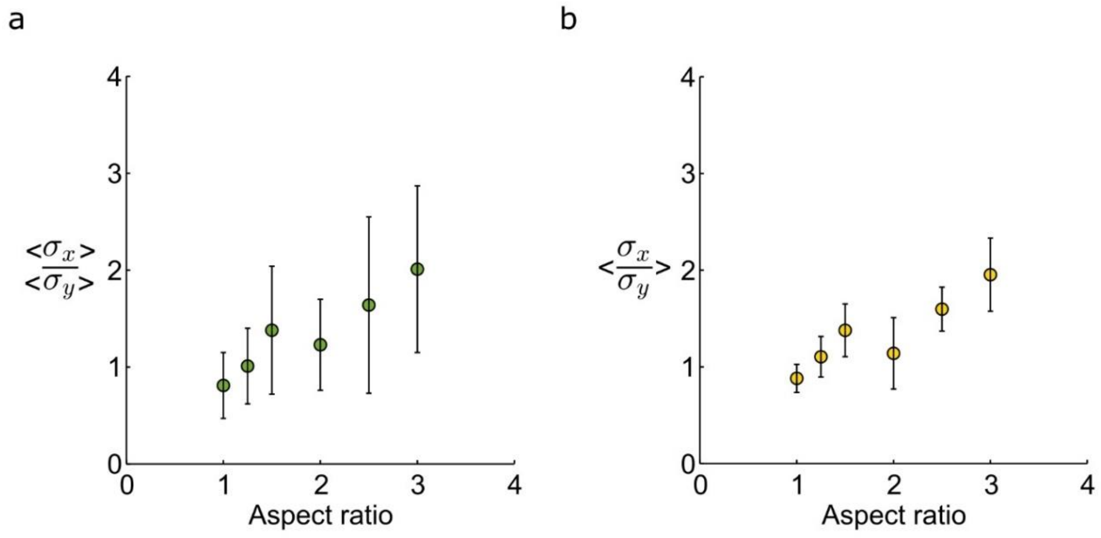


**Figure 3:** (a) Ratios of the mean values $<\sigma_x>$and $<\sigma_y>$ as function of the cell aspect ratio. The ratio seems to be increasing with the cell's aspect ratio, but data are quite noisy with a high SEM preventing one from a clear conclusion. (b) Mean of the ratios $\frac{\sigma_x}{\sigma_y}$ measured for each single cell as function of cell aspect ratio. As shown, this ratio measured cell by cell thanks to our bi-axially calibrated micro-rod force-probe reduces the data dispersion due to variability from cell to cell. In these conditions, the increase of $\frac{\sigma_x}{\sigma_y}$ with the cell's aspect ratio appears significant, the surface tension along the x axis getting higher than the surface tension along the y axis with increasing AR. Roughly, from AR= 1 to AR= 3, $\sigma_x$ gets twice as big as $\sigma_y$.

### Supplementary figures

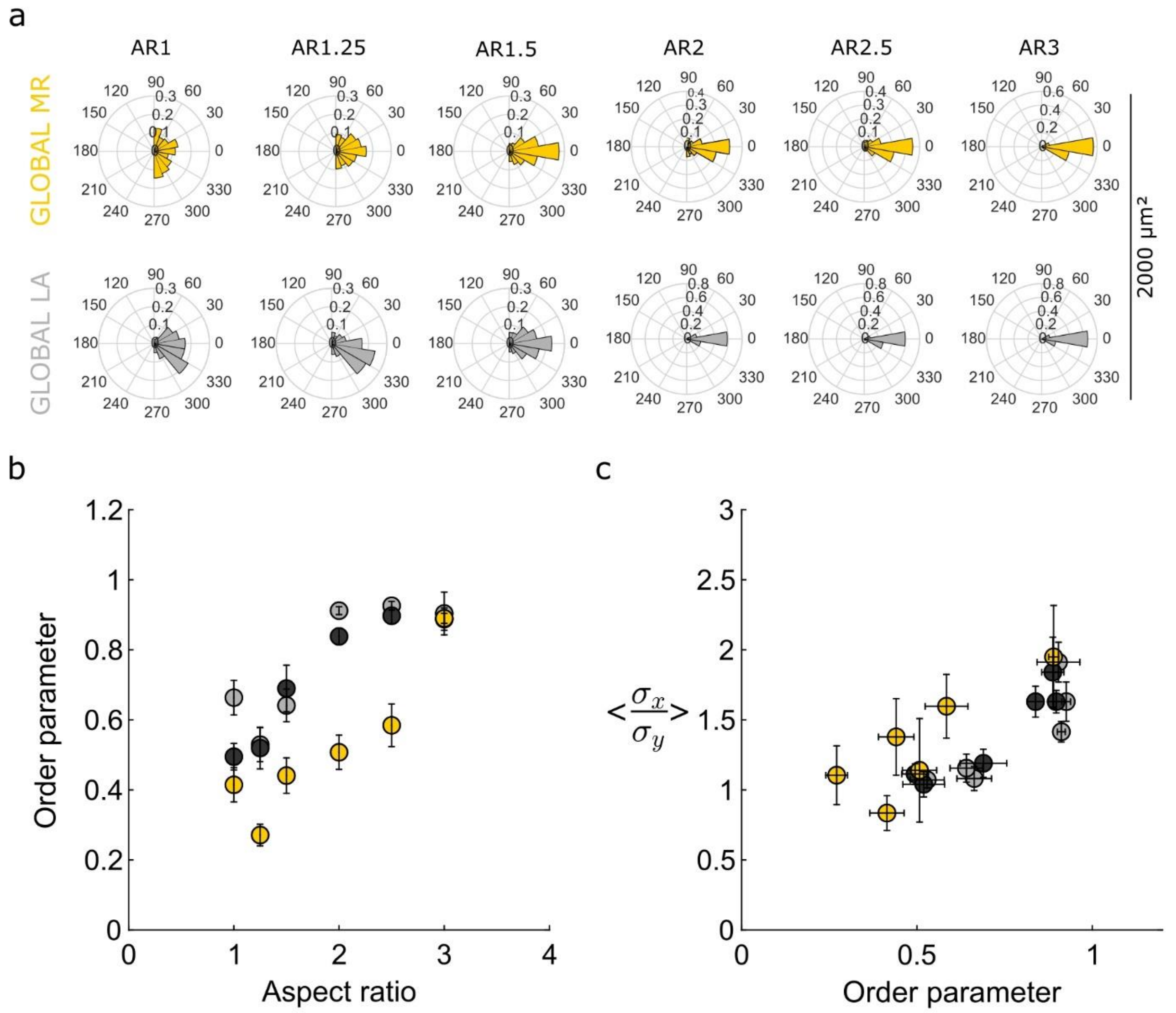


**Figure S1:** (a) Polar histograms of the orientations of the cortical stress fibres (CSF) as observed during line tension measurements (yellow) and laser ablation experiments (grey). (b) Order parameter $S$ as a function of the cell aspect ratio and (c) mean values of the ratio of surface tensions $< \frac{\sigma_x}{\sigma_y} >$ as a function of the order parameter, for line tension measurements (yellow) and laser photoablation experiments (light grey: cells spread over an area of 2000µm²; dark grey: 3400µm²) .

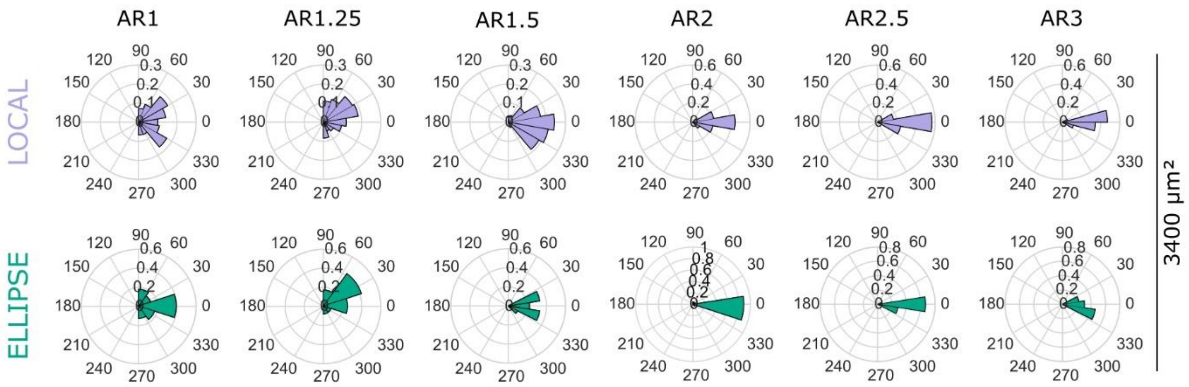


**Figure S2:** Polar histograms of the local orientations of the cortical stress fibres (purple) before photoablation, and that of the ellipses open after laser ablation of the same area (green) obtained for the largest spread area of 3400µm$^2$.
